# Tripeptide loop closure: a detailed study of reconstructions based on Ramachandran distributions

**DOI:** 10.1101/2021.05.23.445336

**Authors:** T. O’Donnell, C. H. Robert, F. Cazals

## Abstract

Tripeptide loop closure (TLC) is a standard procedure to reconstruct protein backbone conformations, by solving a zero dimensional polynomial system yielding up to 16 solutions. In this work, we first show that multiprecision is required in a TLC solver to guarantee the existence and the accuracy of solutions. We then compare solutions yielded by the TLC solver against tripeptides from the Protein Data Bank. We show that these solutions are geometrically diverse (up to 3Å RMSD with respect to the data), and sound in terms of potential energy. Finally, we compare Ramachandran distributions of data and reconstructions for the three amino acids. The distribution of reconstructions in the second angular space (*φ*_2_, *ψ*_2_) stands out, with a rather uniform distribution leaving a central void.

We anticipate that these insights, coupled to our robust implementation in the Structural Bioinformatics Library (https://sbl.inria.fr/doc/Tripeptide_loop_closure-user-manual.html), will boost the interest of TLC for structural modeling in general, and the generation of conformations of flexible loops in particular.

## 1 Introduction

### Conformational diversity of biomolecules

The structure - dynamics - function paradigm stipulates that it is the structure and dynamics of biomolecules which account for their function. Molecular flexibility in the realm of molecular mechanics encompasses vastly different time scales (from picoseconds to seconds) and amplitudes (from milliangstroms to angstroms) [1]. A convenient framework to think about these is that of energy landscapes [2], which decouples structure (identifying meta-stable states), thermodynamics (computing statistical weights of such states), and dynamics/kinetics (modeling transitions using say Markov state models). While flexibility covers very different scenarios, two prototypical ones are of special interest for globular proteins, as they involve loops. In the first, one, which may be ascribed to structural changes, flexibility drives large amplitude conformation changes between meta-stable states involving rigid domains connected by linkers [3], a process which is key for enzymatic function [4] or the efflux by complex membrane proteins [5], to take two examples. In the second one, which may be ascribed to thermodynamics, more local fluctuations of loops contribute to statistical weights whence free energies, a classical implication being an enhanced binding affinity due to a lesser entropic penalty upon binding for pre-structured loops [6]. A different realm is that of intrinsically disordered proteins (IDPs), whose structural plasticity is often linked to biological functions and diseases [7]. IDPs exist as an ensemble of rapidly interconverting structures defining plateaus on the free energy landscape as opposed to the wells associated with stable structures [8]. While differences with globular proteins in terms of Ramachandran distributions have been characterized [9], predicting IDPs properties remains a challenge, and there has been recent awareness of the need for force field modifications (e.g. [10]).

Predicting conformational changes for loops is in fact a hard problem, be it restricted to structure [11] or thermodynamics [6]. A core difficulty for such prediction methods is the inherent bias imposed by the datasets, extracted from the Protein Data Bank, used to calibrate general methods. By construction, experimentally resolved structures incur a bias towards stable structures, so that transient conformations are not accessible. We note in passing that in the aforementioned framework of energy landscapes, transient conformations are generally associated with saddle point regions on the potential energy surface, namely points whose identification requires numerical procedures [12].

### Generating conformations and the Tripeptide Loop Closure problem

Generating diverse confor-mations requires sampling the conformational space. Because dihedral angles are in general softer than bond lengths and valence angles, methods of choice are those restricting the sampling to the former. Narrowing down the focus further, the tripeptide loop closure problem (TLC) consider the six dihedral angles *φ, ψ* found apart from three consecutive *C_α_* carbons. The TLC problem has a long history in robotics and molecular modeling, see e.g. [13, 14, 15, 16, 17, 18]. Mathematically, consider a tripeptide whose internal coordinates (bond lengths {*d_i_*}, valence angles {*θ_i_*}, and dihedral angles {*φ_i_, ψ_i_, ω_i_*}) have been extracted. The TLC problem consists of finding all geometries of the tripeptide backbone compatible with the internal coordinate values {*d_i_, θ_i_*}. Solving the problem requires finding the real roots of a degree 16 polynomial, which also means that up to 16 solutions may be found [16, 19].

Having mentioned above the difficulty of predicting conformations for structurally diverse loops [11], we note that tripeptides recently proved instrumental for this endeavor [20]. In a nutshell, the method grows the two sides of a loop by greedily concatenating (perturbed) tripeptide geometries to the chains being elongated, and closes the loop by solving a TLC problem. Two key steps of the method are the perturbation and sampling from a database of tripeptides used (derived from SCOP), and the final TLC step.

### Ramachandran distributions

The TLC problem is also closely related to the study of Ramachandran distributions, which characterize the coupling between *φ* and *ψ* angles along the protein backbone [21]. There are four main types of Ramachandran plots: glycine – an amino acid without side chain, proline – whose cycle induces specific constraints, pre-proline – residues preceding a proline, and the remaining amino acids, whose *C_β_* carbon induces specific constraints. In this work, we illustrate this latter class with ASP. Four main regions are occupied in the Ramachandran diagram: *β*-sheets (*βS*), polyproline II (*βP*; left-handed helical structure whose angles are characteristic of *β*-strands); *α*-helical (*αR*); and left handed helix (*αL*). These regions were characterized using a combination of five steric constraints between four atoms defining the Ramachandran tetrahedron ([22], Fig. 4). (We note in passing that the 6th edge of this tetrahedron, between *O_i_* and *N_i_i* + 1, was not used in defining the steric constraints, likely due to the fact that this edge corresponds to a valence angle – a constraint stronger than that associated with the other edges.) In this work, the curves delimiting the occupied regions are termed the Ramachandran *template*. More recently the diagonal shape of level set curves in the occupied regions was explained using dipole-dipole interactions, distinguishing the generic case and proline [23], and glycine and pre-proline [24]. The characterization of neighbor dependent Ramachandran distributions has also been studied [25]. From a statistical standpoint, the Ramachandran distributions of two specific residues can be compared using say *f*-divergences such as Kullback-Leibler, Hellinger, etc.

**Figure 1:**
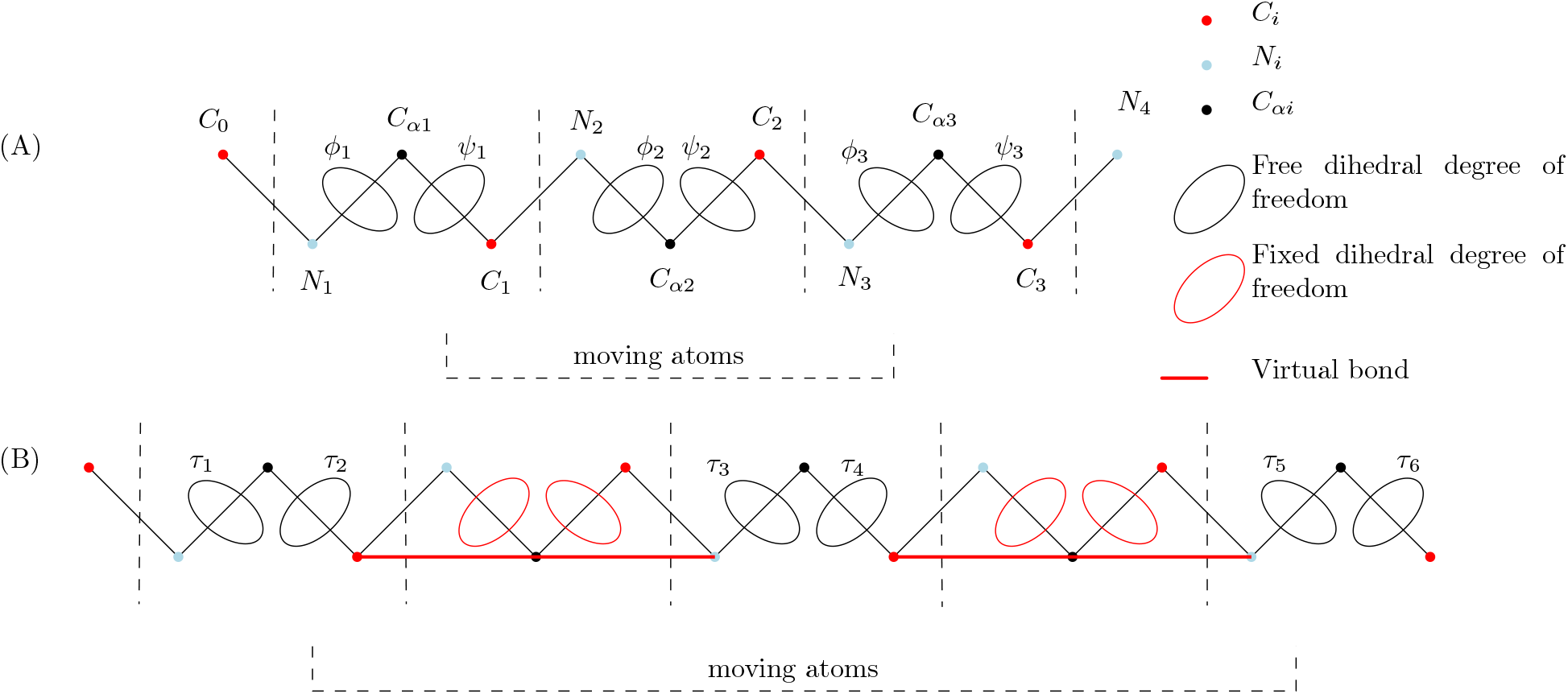
Tripeptide: atoms and degrees of freedom used for loop closure. **(A)** Classical tripeptide loop closure(TLC): the six dihedral angles represented correspond to the degrees of freedom used to solve the problem. **(B)** In tripeptide loop closure with gaps(TLCG), the dihedral degrees of freedom *τ_i_* may be separated from each other by gaps.

**Figure 2:**
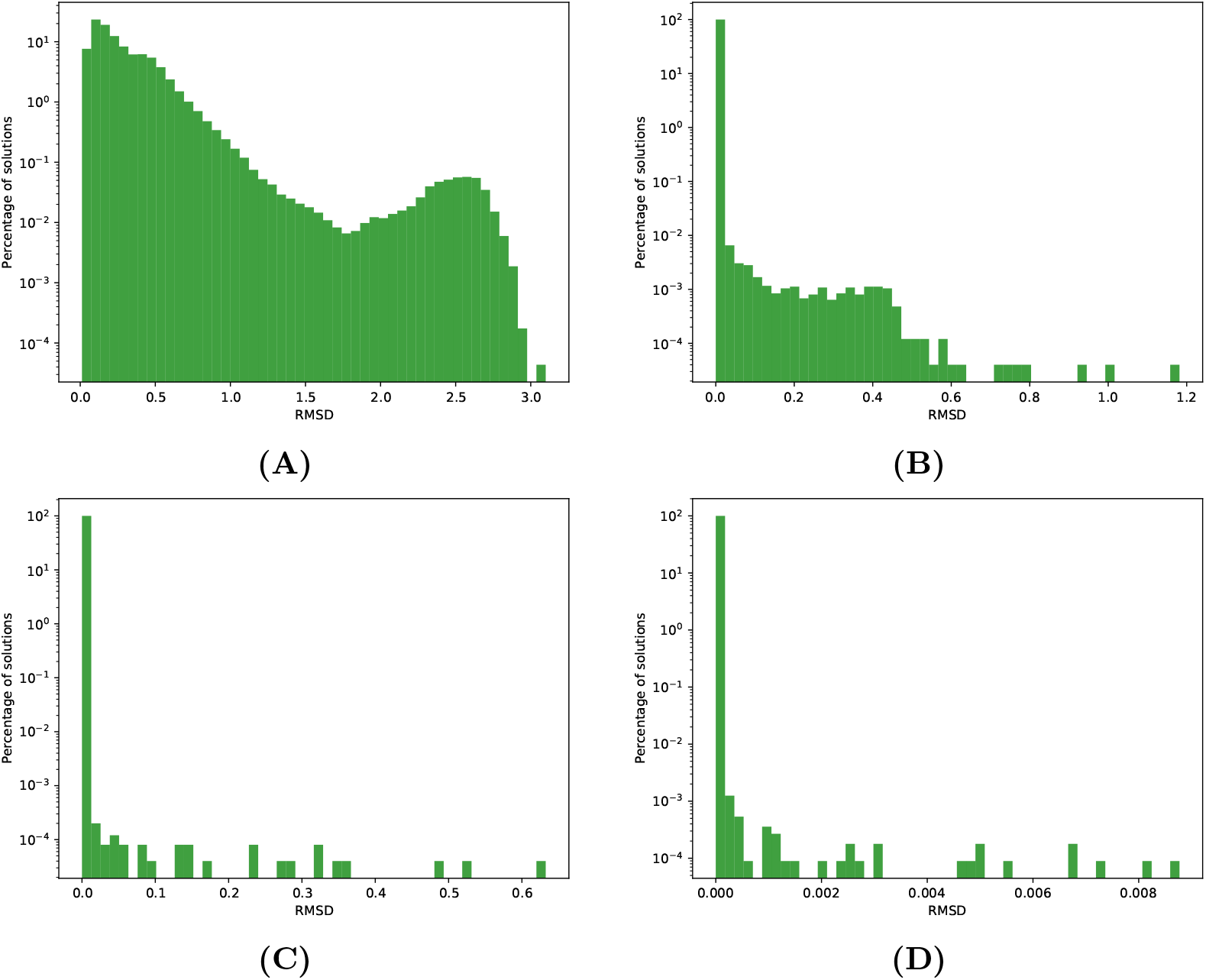
Minimum RMSD between the reconstruction geometrically most similar (RMSD in Å) to the associated data tripeptide. **(A)** `TLCCoutsias` **(B)** `TLCdouble` **(C)** `TLCdouble`[-x2] – twice precision in mantissa **(D)** `TLCdouble`[-x4] – quadrice precision in mantissa

**Figure 3:**
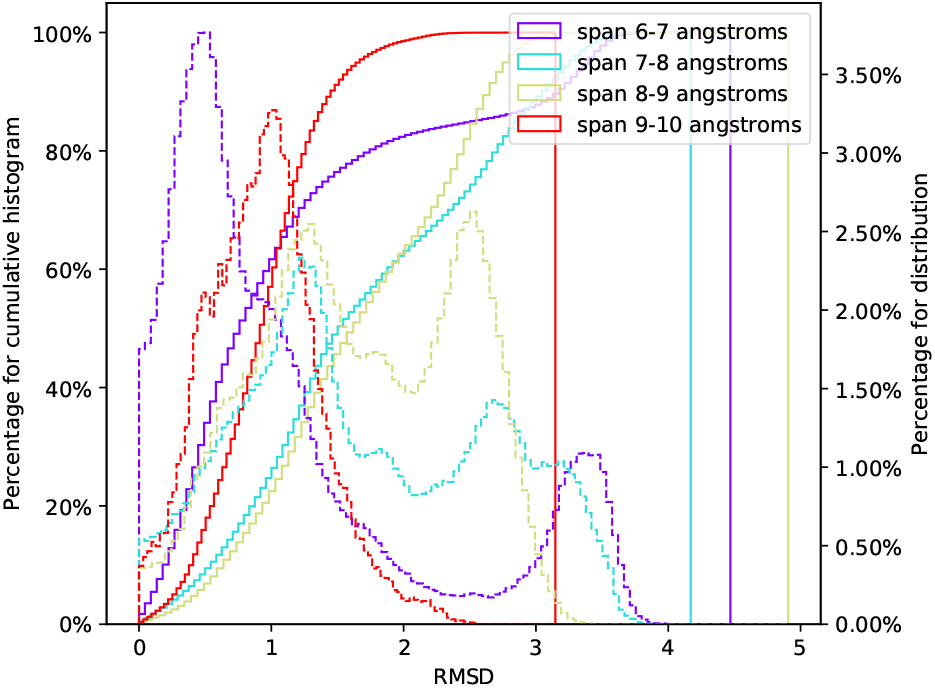
Solutions yielded by TLCdouble[-x2]: maximum RMSD between each set of reconstructions and the original data. Upon solving *TLC*(*l*) for a tripeptide *l*, the solution most dissimilar to *l* in the RMSD sense is sought in the solution set *Sol*(*l*) = {*r*_1_*, …, r_k_*}. The full line represents the cumulative histogram of this maximal RMSD, with the corresponding Y axis on the left. The dashed line is a regular histogram of the same data, with the corresponding Y axis on the right.

**Figure 4:**
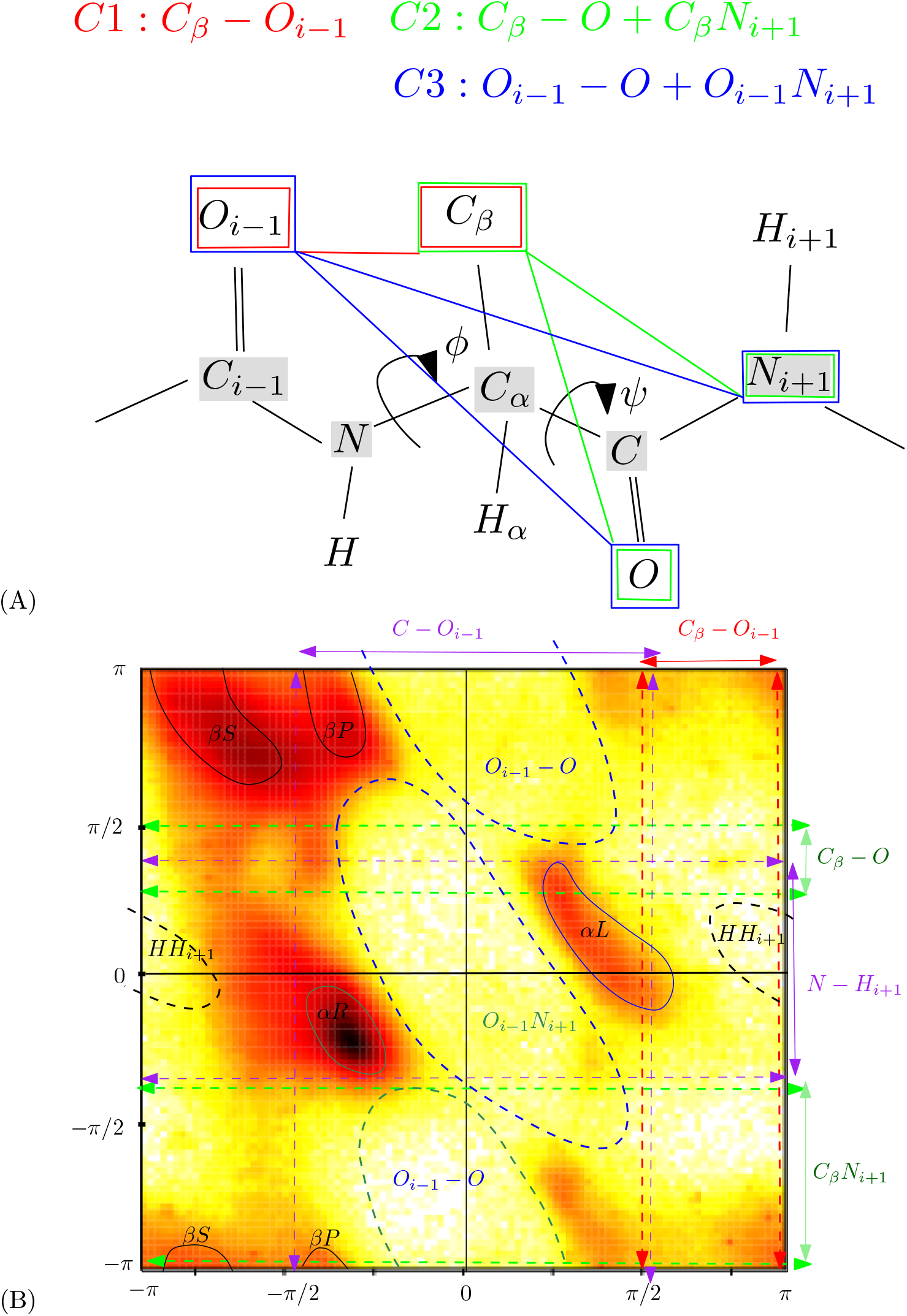
Ramachandran diagrams: distance constraints and occupied regions. **(A)** The Ramachandran *tetrahedron* and its five distance constraints – adapted from [23, 24]. Note that the four atoms define a tetrahedron: five of its edges are constrained; the last one (*ON_i+1_*) corresponds to a valence angle, and is not constrained. **(B)** Main regions occupied in the Ramachandran space, with associated steric constraints, materialized by dashed lines/curves, involving vertices of the Ramachandran tetrahedron. The background distribution was obtained using all amino acids in the structure files used in this study (loops and SSE). The partition of the Ramachandran space illustrates the location of the classical SSE: *β*-sheets (*βS*), polyproline II (*βP*; a left-handed helical structure whose angles are characteristic of *β*-strands), *α*-helical (*αR*), and left handed helix *αL*.

### Contributions

In this work, we perform a careful assessment of reconstructions to TLC problems, with a particular emphasis on the comparison between distributions in angular spaces, between data from the PDB on the one hand, and TLC reconstructions on the other hand.

First, we present a robust implementation of TLC, showing the role of multiprecision in ensuring the existence and the accuracy of reconstructions. Second, using tripeptides from the PDB as a reference, we present a detailed analysis of reconstruction, from the geometric, statistical, and biophysical standpoints. We also discuss some possibilities to exploit such reconstructions.

## 2 Material and Methods

### 2.1 Material: tripeptides from the PDB

We extract a database 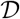 of tripeptides found in high resolution structures (resolution better than 3Å) from the PDB (23rd of September 2020), having mutual sequence identity lower than 95%. A contiguous, gap-less region of a protein backbone qualifies as a tripeptide if the following two conditions hold: (C1) The highest Bfactor in backbone atoms of the tripeptide is less than 80 Å^2^. (C2) The center of the tripeptide is separated by at least 3 amino acids from a stable secondary structure(SSE) on both ends, a condition meant to remove the constraint of SSE anchoring loops to the rest or the structure [25]. Stable secondary structure (SSE, *β* folds and right handed *α* helices) are extracted from mmtf files. These files are annotated using the BioJava implementation [26] of the DSSP program (Define Secondary Structure of Proteins [27]).

In order to compute the original values of the first *φ* and last *ψ* dihedral angles, the tripeptide at the end or beginning of a chain is excluded from our computation as the positions of the last atom of the previous residue and the first of the next one are necessary. Taken together these conditions result in the database 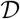 containing 2,495,095 tripeptides. We denote 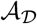 the corresponding encoding in the 6D space of dihedral angles, that is

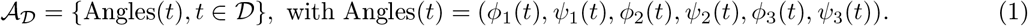

We qualify a tripeptide with its span (Euclidean distance between its endpoints the *N*_1_ and *C*_3_ atoms (Fig. S1). In computing the percentage of tripeptides containing a given amino acid at least once, Glycine is followed by Proline and Aspartic acid (29.6%, 24.7%, 21.6% of tripeptides respectively) (Table S1(A)). For the percentage of tripeptides containing a particular amino acid at least twice, this ordering remains the same, the relative gap between glycine and the following amino acids being wider (Table S1(B)).

### 2.2 The classical TLC problem

#### Data versus reconstructions

We consider the tripeptide loop closure with fixed bond lengths and angles, as well as *ω* dihedral angles. As mentioned previously both sides of this tripeptide are fixed (i.e. *C_O_, N*_1_*, C_α_*1 and *C_α_3, C*_3_*, N*_4_), meaning that the collective change in dihedral angles only affect the Cartesian embeddings of *C*_1_, *N*_2_, *CA*_2_, *C*_2_ and *N*_3_ (Fig. 1(A)). For a TLC problem defined by a tripeptide *t*, the set of solutions and the ancestor of a solution are denoted as follows (Fig. S1 for one example):

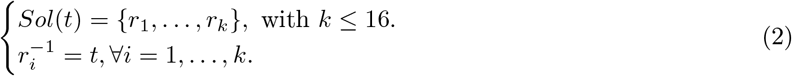

A TLC problem is expected to return the tripeptide it is defined from. (As we will see, this depends on the number type used.) In the solution set *Sol*(*t*), we will therefore assume that *r*_1_ is the reconstruction most similar to the data tripeptide *t*, in the RMSD in 3D space sense. (Setting aside numerical precision issues, the data tripeptide should be exactly reconstructed, i.e. the RMSD should be zero.) We define accordingly the solution set minus the data tripeptide, that is

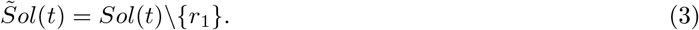

#### Angular spaces

Denote *d_S_*1 (·, ·) the shortest distance between two angles on the unit circle *S*_1_. To compare tripeptides whose 6D dihedral coordinates are denoted Angles(*t*) = (τ_1_, …, τ_6_) and 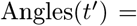)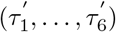 respectively, we use as distance the *L_p_*-norm – in practice with *p* = 1:

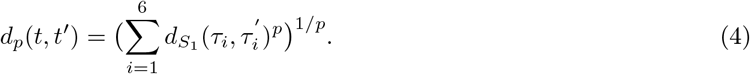

We also consider the following angular data associated with all reconstructions:

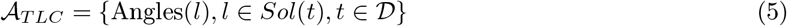

With the specific goal of analyzing reconstruction which differ from the original data, we define the subset of solutions

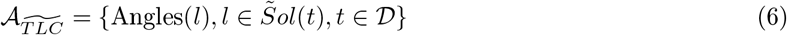

For data in 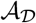 (resp. reconstructions in 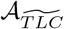), the pairs of dihedral angles of the i-th tripeptide are denoted (*φ_i_, ψ_i_*) 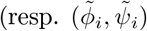) and the corresponding Ramachandran domain is denoted 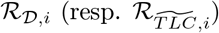.

#### TLC and internal coordinates

Solving a particular TLC problem puts the focus on dihedral angles, so that that there are two options to handle the other internal coordinates (bond length and valence angles): *data internals* using those found in the tripeptide processed, and *canonical internals* using standard values for fixed internals, as done in the original version[19]. As we shall see, the former is beneficial in several respects.

### 2.3 TLC with gaps

A generalization of the classical TLC consists of considering three amino acid which are not contiguous along the backbone. This is of interest in the case of three linkers enclosing two rigid SSE. Mathematically, this is akin to the original problem, with the rigid blocks modeled as fictitious bonds separating the amino acid (Fig. 1(B)). Once the coordinates of all atoms not in these rigid blocks are embedded, the rigid blocks are then translated and rotated into their final positions (Fig. S2 for one example).

## 3 Results

### 3.1 Software

#### TLCG algorithm

This work is accompanied by our implementation of the tripeptide loop closure algorithm, in the Structural Bioinformatics Library ([28], http://sbl.inria.fr, https://sbl.inria.fr/doc/Tripeptide_loop_closure-user-manual.html). From the application standpoint, given a chain in a PDB file, together with the identification of the three a.a. defining the tripeptide (not necessarily contiguous), the application `sbl-tripeptide-loop-closure.exe` produces modified PDB-format files for each solution found, if any. The constraints for internal coordinates can be specified from the data (default), using standard values, or supplied in the form of a file.

#### Numerics

The numerical stability of an algorithm is key to its robustness [29]. For TLC, the precision used to represent the floating point numbers is expected to play a role.

The application sbl-tripeptide-loop-closure.exe makes it possible to specify the precision used for calculations. Internally, the number type used is `CGAL::Gmpfr`, a representation based on the `Mpfr` library [30] supplying a fixed precision floating point number type. Practically, this fixed precision is a multiple (> 1) of the default double precision: `TLCdouble`[−x1], called `TLCdouble` for short in this work, refers to the executable `sbl-tripeptide-loop-closure.exe` using the plain double precision; `TLCdouble`[−x2] (resp. `TLCdouble`[−x4]) refers to `sbl-tripeptide-loop-closure.exe` using a double double (resp. quadrice double) precision.

To assess the importance of using data-extracted (as opposed to standard) internal coordinates, we also evaluate `TLCCoutsias` [19], the original TLC algorithm using standard bond length and valence angles, with double precision for numerics.

### 3.2 Numerical analysis of the stability of the reconstruction

#### Rationale

Solving a TLC problem for a tripeptide *l* raises two questions. The first question refers to the existence of a solution matching the data *l* itself. The response can be negative since numerical rounding errors during the calculation of the polynomial may yield, in particular for an ill-conditioned TLC polynomial, a situation with zero real solution [29]. In that case, we will say that the solution *evaporates*. If solutions are found, we define *the reconstruction* as the geometry most similar to *l*, using as distance the RMSD of the atoms in the tripeptide. Note that RMSD and not least-RMSD is used for this comparison as the orientations are fixed.

The second question is then the geometric distance between the data and the reconstruction. This distance, also measured by the RMSD, is expected to depend on the floating point number type used.

#### Results

We process all cases in the database 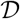. The fraction of TLC problems with no solution depends heavily on the option used for number types and internal coordinates other than dihedrals: `TLCCoutsias`: 8.1%; `TLCdouble`: 5 10^−4^%; `TLCdouble`[−x2]: 2 10^−5^%; `TLCdouble`[−x4]: 0.0. A similar conclusion holds for the RMSD between the data and the (best) reconstruction (Fig. 2(A, B)): `TLCCoutsias`: up to 3Å RMSD; `TLCdouble`: up to 1.2Å RMSD; `TLCdouble`[−x2]: very small values with one outlier at ~ 0.65Å RMSD; `TLCdouble`[−x4]: all RMSDs smaller than ~ 0.009Å RMSD.

Altogether, these observations stress the importance of using data-extracted internal coordinates, and to a lesser extent the role of numerical precision to avoid evaporation. While `TLCdouble` is sufficient to characterize distributions, `TLCdouble`[−x2] is preferable to process satisfactorily all individual cases. In the sequel, all results presented were obtained with `TLCdouble`[−x2].

### 3.3 Geometric analysis of solutions in 3D

#### Rationale

To assess solutions, we consider the reconstruction from 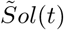 most dissimilar to *t*, in the RMSD sense.

#### Results

For data extracted internals, the number of solutions of TLC problems is as high as 12 (Fig. S3). The analysis of the geometric diversity in terms of max RMSD as a function of the geometric span of the tripeptide (Euclidean distance between its endpoints) yields two interesting insights (Fig. 3). First, with only 5 displaced atoms, a significant RMSD is observed, up to 3.8Å. Second, the distribution is bimodal, but the two modes get closer (and even coalesce) when the span increases. This can be explained by the fact that the larger the gap, the straighter the solutions.

As a complementary analysis, consider the displacement of the 5 moving atoms in the tripeptide (Fig. 1). For each atom, we compare the generated position against the initial position. As expected, the displacement increases with the centrality of the atom, with displacements which can be very significant, namely up to 6 Å(Fig. S4).

### 3.4 Geometric analysis of solutions in 6D

#### Rationale

We wish to perform a geometric comparison of the two 6D point clouds 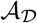 and 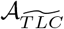 coding all tripeptides in 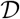 and in all solutions (minus the data tripeptides themselves) respectively (Section 2.2).

To see how, consider two set of points in 6D, say *X* and *Y*. For a point *x ∈ X*, using the distance from Eq. 4, we define the nearest neighbor in *Y* and the associated distance by

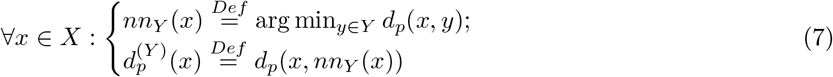

#### Remark 1

*We may need to restrict the search of the nearest neighbor of of a tripeptide x to a certain class tripeptides sharing a specific property with x – e.g. featuring a C_β_. The corresponding operator is denoted 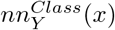.*

#### Results

The distribution of 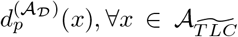 has a sharp mode at zero, showing that ~ 20% of solutions (data tripeptides excluded) are highly similar to a tripeptide existing in 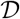 (Fig. S5(A)). Taking the reverse point of view, the distribution of 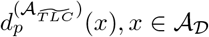 shows that the number of data tripeptides *p* D similar to a solution is ~50% (Fig. S5(B)). Interestingly, the span of values in these two histograms are circa 130 and 40 degrees respectively, showing that loop closure tripeptides are far more diverse than PDB peptides.

### 3.5 Analysis of Ramachandran distributions

#### Rationale

We complement the previous geometric analysis by studying the distributions in Ramachandran spaces. The focus in doing so is twofold: first, comparing the distributions in 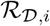 versus 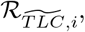, and second, analyzing the patterns observed with respect to those known for classical Ramachandran plots (Fig. 4).

#### Results

We inspect individual Ramachandran distributions over the domains 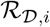 versus 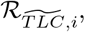, considering three prototypical amino acids, namely ASP (Fig. S6), GLY (Fig. S7), and PRO (Fig. S8). Ramachandran distributions in the domains 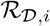(left columns) are indistinguishable (also confirmed by the calculation of the Hellinger and Jensen-Shannon divergences, data not shown), a fact which is expected since tripeptides from the database 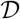 are obtained by sliding a window of size three along loops found in structures from the PDB.

On the other hand, the three Ramachandran distributions associated with the TLC domains 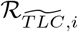 = 1, 2, 3 are rather different. Distributions in the domains 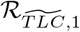, and 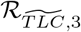 still exhibit isolated regions corresponding to classical regions, except that the distributions are much more uniform in the entire Ramachandran space. The middle distribution (space 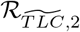) departs from these. The coverage of the entire space is more uniform, setting aside a central void surrounded by an *annulus* connecting the clusters corresponding to the classical structures (left and right handed *α* helices, *β* folds). The central void/eye corresponds to the steric constraint *O_i−1_N_i+1_* (Fig. 4). It should be noticed, though, that the clear cut nature of this void results from the fact data have been removed from the solutions set (Eq. 6). In plotting all pairs of angles (*φ, ψ*) from our database 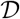, one indeed obtains atypical conformations in this central region (background of Fig. 4). In any case, the superposition of the Ramachandran template onto the map shows that solutions partly fill the void (Fig. 5). Interestingly, the center of the void is preserved even though the distance constraints encoded in the Ramachandran tetrahedron are not used in the specification of the TLC problem – since *N_i+1_* only is involved in the TLC problem (Fig. 4). The comparison of maps in 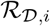 and 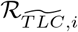 is provided by the difference plots (Figs. S6, S7, and S8). In addition to stressing the differences already mentioned, we note that, in accordance with the steric constraints found before and after the tripeptide, the first (resp. third) difference plot exhibits a vertical (resp. horizontal) stripe, showing that *φ*_1_ is more constrained than *ψ*_1_ (resp. *ψ*_3_ more constrained than *φ*_3_).

**Figure 5:**
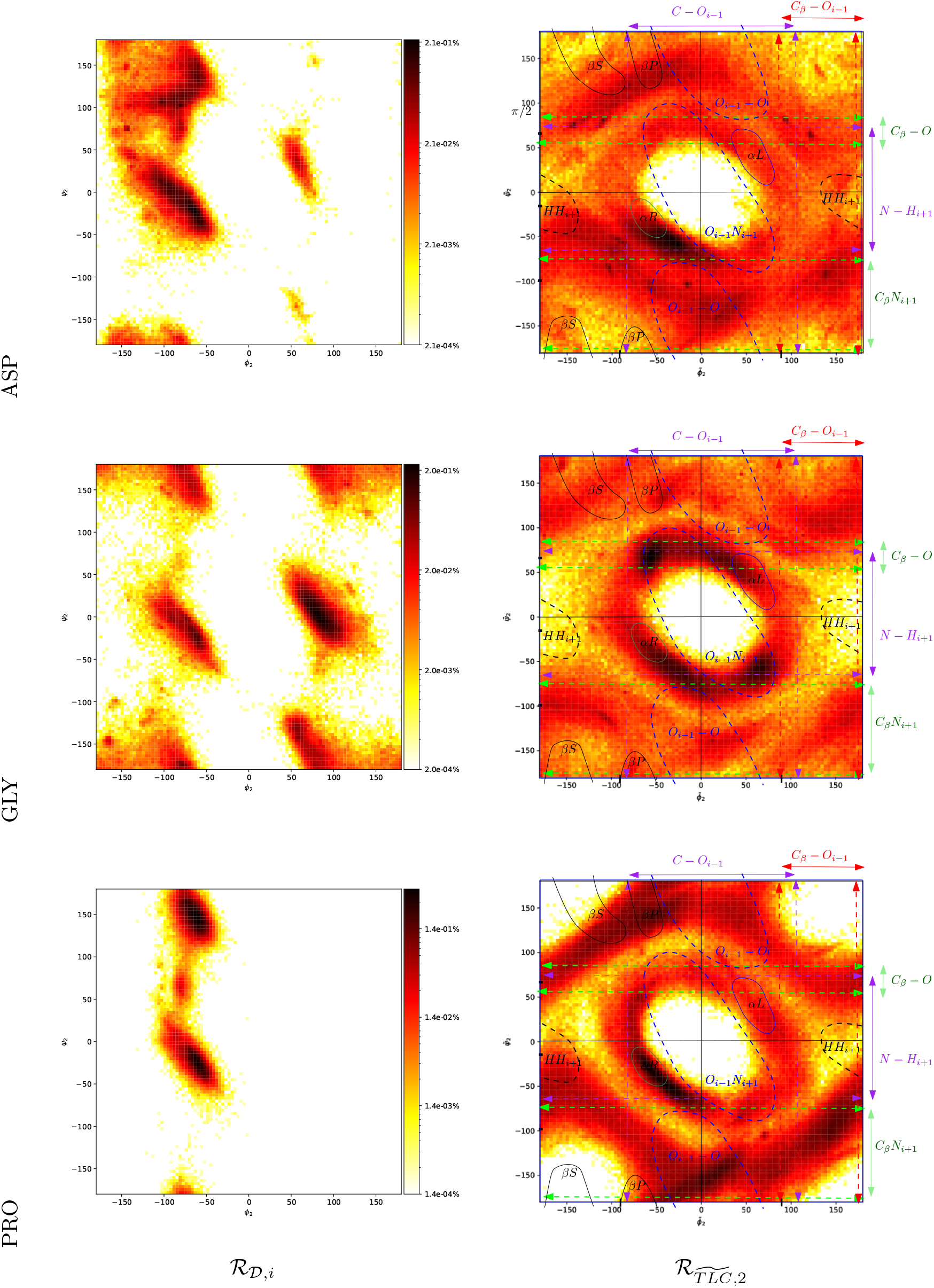
Ramachandran distributions for ASP, GLY, and PRO. **(Left column)** Distributions for domains 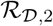 **(Right column)** Distributions for domains 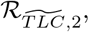, with the superimposed Ramachandran template.

### 3.6 Biophysical analysis based on the potential energy of solutions

#### Rationale

As a separate assessment of the quality of reconstructions returned by `TLCdouble`, we compute the potential energy (denoted *V*) of the tripeptide backbone including heavy atoms (*i.e.* the carbonyl oxygens and *C_β_*) involved in the specification of the regions occupied in Ramachandran diagrams (Fig. 4). This analysis imposes two constraints. First, we discard tripeptides containing PRO. Second, we assign a type to each tripeptide, out of 2^3^ possibilities corresponding to the presence or absence of a GLY at each position. This type is used in particular to find the nearest neighbor of a reconstruction amongst all tripeptides of the same type in 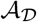, which we denote 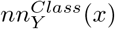. (See Rmk 1. In Eq. (7), the set *Y* is filtered to retain those tripeptides whose type matches that of *x*.) Practically, we present plots for the most abundant class, corresponding to tripeptides with a *C_β_* at each position.

Three potential energy terms are taken into account:

- The first corresponds to the contribution of dihedral angles. Each such angle contributes Σ*_n_*(*k*(1 +cos(*nφ-φ_0_*))) with *n* the periodicity of the term, *k* the energy constant, *φ*_0_ a phase shift angle, and *φ* is the torsion angle formed by the four bonded particles.
- The second term is the electrostatic interaction between non bonded particles. Each non bonded pair contributes 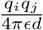, where *E* is the dielectric constant, *q_i_, q_j_* are the charges of the two particles, and *d* is their distance.
- The last term is the van der Waals interaction term. Each non bonded pair contributes 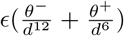 where *E* is a constant, *θ*^−^*, θ*^+^ are the repulsive and attractive Lennard-Jones terms, and *d* is the distance between particles.

In any case, only contributions impacted by the changes made by the TLC algorithm are taken into account. For the dihedral angles, this implies that proper dihedrals around the peptide bonds are not taken into account. For the non bonded interactions only pairs whose relative distance changes contribute.

To assess the potential energy of a reconstruction in 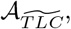, we compare this potential energy to a reference point. This can either be the nearest neighbor of each point 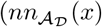, Eq. 7) or the data *x*^−1^ used to generate it (Eq. 2). This yields the following two relative changes for the potential energy *V* (·) with * {*dihedral, elec., vdW*}:

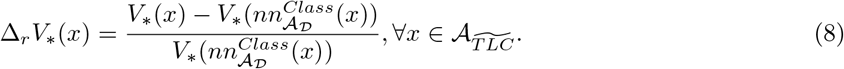

or

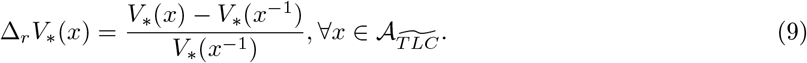

Using the whole database, we perform a scatter plot in the plane 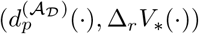, and represent the resulting 3D histogram using a heatmap.

#### Results

The potential energy is a measure of the *strain* of reconstructions. The analysis of the three potential energies and the two comparison setups yields several interesting facts (Fig. 6):

**Figure 6:**
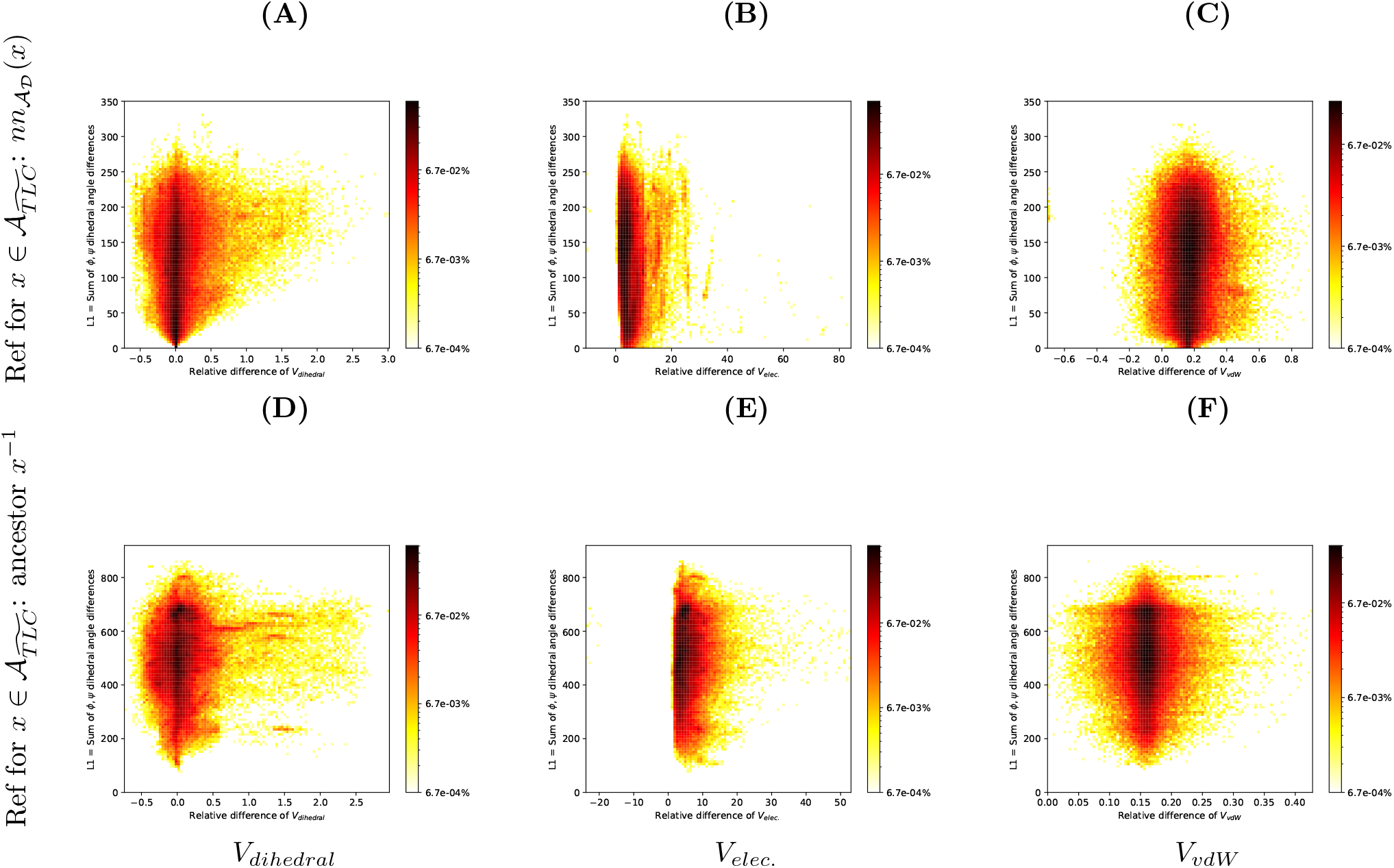
Relative changes of the potential energy: reconstructions in 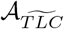 versus a reference tripeptide, for all tripeptides of class ASP (i.e., without GLY and featuring a *C_β_* at each position). Calculations involve all backbone heavy atoms, including the carbonyl oxygen and the *Cβ*. The y-coordinate is the sum of angular distances to the match used (L1 norm, Eq. 4). The color depends logarithmically on the percentage of all solutions in a bin. **(Top row)** Reference tripeptide for a reconstruction 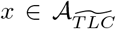 is the nearest neighbor of the same class 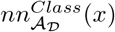. **(Bottom row)** Reference tripeptide *x*^−1^ for a reconstruction 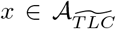 is the ancestor *x*^−1^ **(First column)** Potential energy of dihedral angles. **(Second column)**Electrostatic term, involving all pairs of atoms whose relative distance changes. **(Third column)** van der Waals term, involving all pairs of atoms whose relative distance changes.

- **Magnitude of angular changes.** We note that using the nearest neighbor of a reconstruction significantly reduces the L1 distance (Fig. 6: from [100, 750] to [0, 250] degrees), an indication on how a reconstruction from 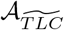 differs from its data tripeptide in terms of dihedral angles.
- **Magnitude of potential energy changes Δ*V*_*_ in kcal/mol – Table S2.** The absolute difference Δ*V*_∗_ has a different scale for the three potential energy terms used: *V_dihedral_* yields the smallest changes, then *V_elec._* and finally *V_vdW_*. The low energetic impact of changes to dihedral angles is what makes it a priority target to modify structures in protein molecules and why TLC is such an interesting approach. The changes in *V_elec._* in the backbone of proteins are more sensitive to modifications done by TLC as the energy linearly depends on the inverse of the distance *d* between non bonded atoms. In the same spirit, with a larger exponent (*d*^12^), the changes in *V_vdW_* are the largest ones. It should be noted that *V_vdW_* (*x*^−1^) has a larger value than the difference Δ*V_vdW_* (*x*).
- **Magnitude of relative changes – Table S3)**. Out of the three potential energies, *V_vdW_* displays a significant difference in terms of relative changes for the two reference tripeptide definitions: from [−0.1, 0.5] (Fig. 6(C)) to [0.05, 0.25] (Fig. 6(F)). Even though a reconstruction resembles less its ancestor than its nearest neighbor in terms of angular coordinates, the spread of relative changes is smaller for ancestors.
- **Centering and symmetry of relative changes.** Relative changes for *V_dihedral_* exhibit a relative symmetry about Δ*_r_V_dihedral_* = 0 (Fig. 6(A,D)), which is expected due to the periodic form of this potential energy. A relative symmetry is also observed for *V_vdW_*, about Δ*_r_V_vdW_* ∼ 0.17 and Δ*_r_V_vdW_* ∼ 0.16 respectively (Figs. 6(C,F)). This negative value shows that data tend to have a smaller *V_vdW_*, yet reconstructions occasionally yield more favorable interactions. Finally, *V_elec._* only displays negative values for relative changes (Figs. 6(B,E)), stressing the rather tight optimization of this potential energy in native structures.
- **Distance** *d*_1_ = 0 **does not imply** Δ*V_dihedral_* = 0. It also appears that *d_p_* → 0 implies Δ*_r_V_dihedral_* → 0 (Fig. 6(A)). This can be explained by considering that if there is no difference in free dihedral angles then the energy term obtained corresponds to that of its reference. This is not true however for Δ*_r_V_vdW_* and Δ*_r_V_elec._*. When using 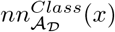 as reference these are impacted by the differences in the other internal coordinates, differenceAs*D*that impact interatomic backbone distances.

## 4 Discussion and outlook

Tripeptide loop closure (TLC) is a classical strategy to generate conformations of tripeptides, e.g. to reconstruct missing segments in structural data, or to implement move sets in simulation methods. Specifically, a TLC problem solves for six dihedral angles, keeping the remaining internal coordinates (bond lengths, valence angles) constant. Solutions are determined by the real roots of a degree 16 polynomial, which makes it very convenient to generate discrete conformations, but which raises questions regarding the biophysical relevance of solutions. The focus of this work is precisely to provide a detailed assessment of reconstructions, using tripeptides from the protein data bank as a reference.

From the computational standpoint, we show that multiprecision is required for the existence and the accuracy of reconstructions. From the geometric standpoint, it appears that the number of solutions depends on the endpoint to endpoint distance of the gap to be filled. Also, despite the fact that a mere five atoms are moving, RMSD up to ~ 6Å are observed, showing that TLC yields a significant conformational diversity. From the statistical standpoint, we present a detailed comparison of angular distribution in the Ramachandran spaces of data and reconstructions, for each of the three positions in the tripeptide. The specific distribution for the second tripeptide in reconstructions is remarkable. This distribution features a central empty region –the *void* pattern, and is more uniform than classical Ramachandran distributions. Such differences actually owe to the different nature of these two distributions. On the one hand, classical Ramachandran distributions encode propensities observed in protein structures, typically extracted from the protein data bank. Such structures are biased toward (meta-)stables states, and one expects transient regions to be under-represented. On the other hand, Ramachandran distributions associated to reconstructions inherently encode the propensities of angles in the TLC reconstructions, which, as we have seen, endow the central atoms of the tripeptide with enhanced move capabilities.

The results thus show that, while reconstructions are themselves conditioned to the input PDB data, their bias towards (meta-)stable structures is less pronounced. Application-wise, the void pattern provides strong hints on how to interpolate between two tripeptides geometries. Given two conformations encoded by two points in the 6D space, one may indeed attempt to connect them while staying away from the void region in a manner akin to path planning in robotics. This strategy remains to be explored, and may be particularly applicable to conformational sampling in less-structured systems such as intrinsically disordered proteins (IDPs), which would accompany recent awareness of the need for force field m odifications. Fi nally, from the biophysical standpoint, we show that the potential energies associated with dihedral angles, electrostatic and van der Waals interactions incur changes of increasing magnitude, in this order. Non bonded distances are not considered in TLC and get impacted more significantly, t he i mportance o f c hanges d epending on the weighting of the interatomic distance (via the distance exponent). Fully assessing the relevance of these solutions requires further work, though. While local steric clashes may arise from the tripeptide geometry provided by solutions of TLC, such clashes may be palliated by performing a local repacking, or by minimizing the overall potential energy, as classically done in methods such as basin-hopping.

Overall, our work furthers our understanding of tripeptide geometries and their link to reconstructions yielded by the tripeptide loop closure. From the software standpoint, we anticipate that our robust open source implementation, available in the Structural Bioinformatics Library will ease the use of TLC in various structural modeling projects in general, and the generation of conformation of flexible loops in particular.

## 5 Artwork

## 6 SI: Methods

### 6.1 Material: loops and tripeptides from the PDB

- Table S1

### 6.2 The TLC geometric model

- Fig. S1
- Fig. S2
- Fig. S3
- Fig. S4
- Fig. S5

### 6.3 Statistical analysis

#### Ramachandran distributions and their difference

For a given a.a. found at position *i* = 1, 2, 3 in a tripeptide, we consider the Ramachandran distribution in spaces 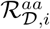 and 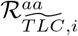 respectively. Furthermore, we define the difference between these distributions in the two dimensional space defined by the signed differences Δ*φ_i_* and Δ*ψ_i_*, as follows:

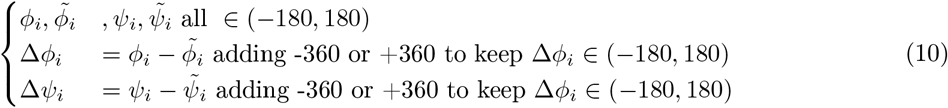

### 6.4 Biophysical analysis

Pairs of atoms contributing to the non bonded terms in eq. 8 and 9: All atoms pairs containing at least one impacted embedding of a heavy atom.

- All pairs containing C1
- All pairs containing O1
- All pairs containing N2
- All pairs containing CA2
- All pairs containing CB2
- All pairs containing C2
- All pairs containing O2
- All pairs containing N3

Any dihedral containing at least one of the atoms above is considered as contributing to the potential energy term relative to the impacted dihedral angles.

---

**Figure S 1:**
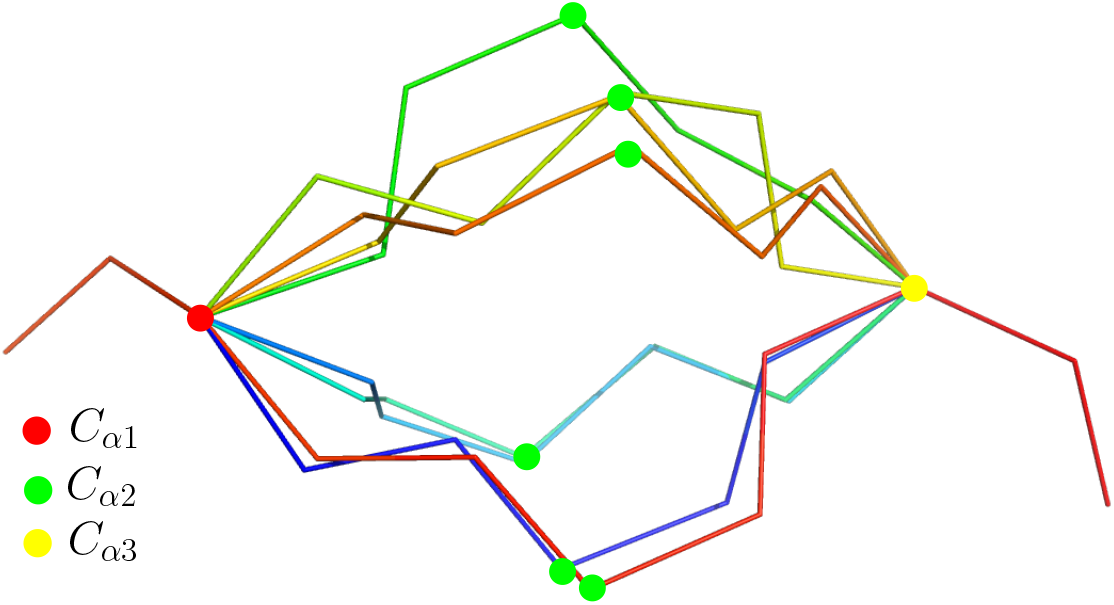
TLC: example reconstructions.

**Figure S 2:**
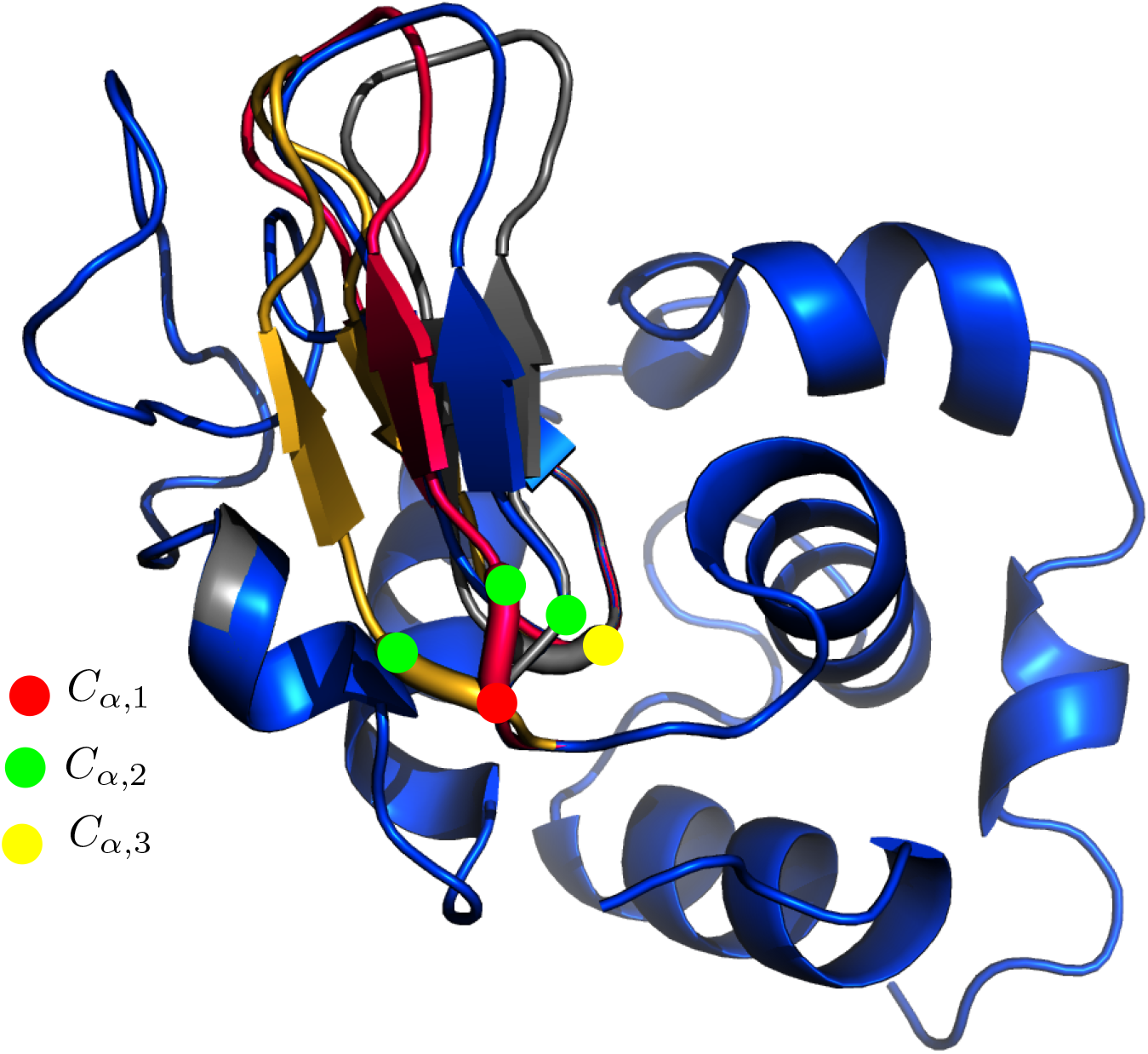
TLCG: example reconstructions sandwiching a beta sheet. PDBID 1vfb, chain C. The three amino acid defining the tripeptide are: *C_α;1_* (resid: 41 GLN), green *C_α;2_* (resid: 42 ALA), yellow *C_α;3_* (resid: 54 GLY). A total of six reconstructions were obtained with TLCdouble[-x2]. Four are displayed for the sake of clarity. The blue one represents the original geometry.

**Figure S 3:**
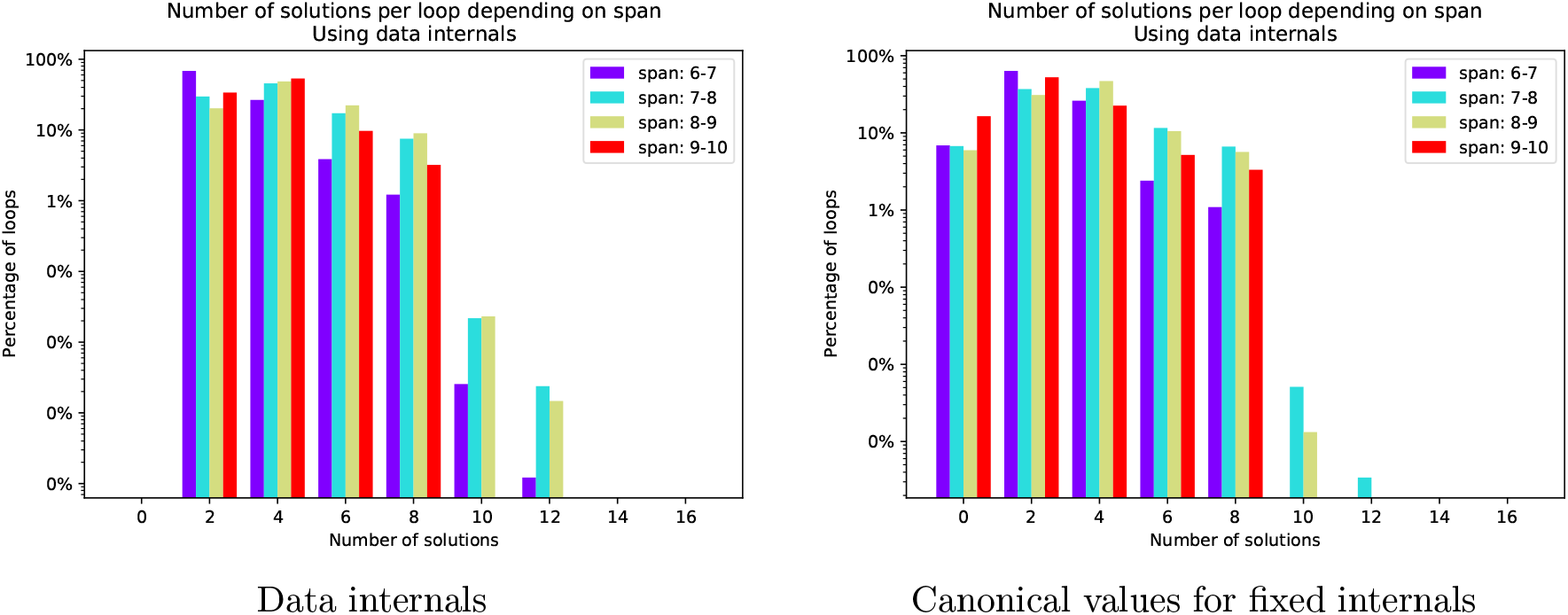
Number of solutions for all TLC problems in our database 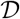. **(Left)** Fixed internals (bond lengths, valence angles) from the data **(Right)** Canonical values for these internal coordinates, from [19].

**Figure S 4:**
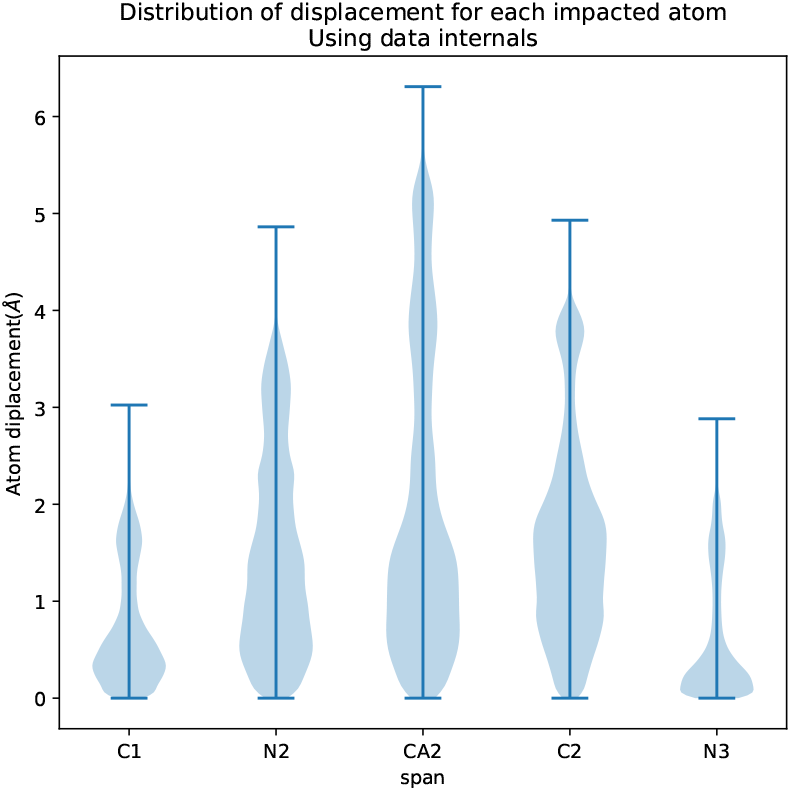
Distribution of displacement for the five moving atoms. Solving a TLC results in five moving atoms (Fig. 1). For all displaced atoms in the loop closure generated solutions this is the distribution of the displacement in Angstroms when compared to the original data used to formulate the loop closure.

**Figure S 5:**
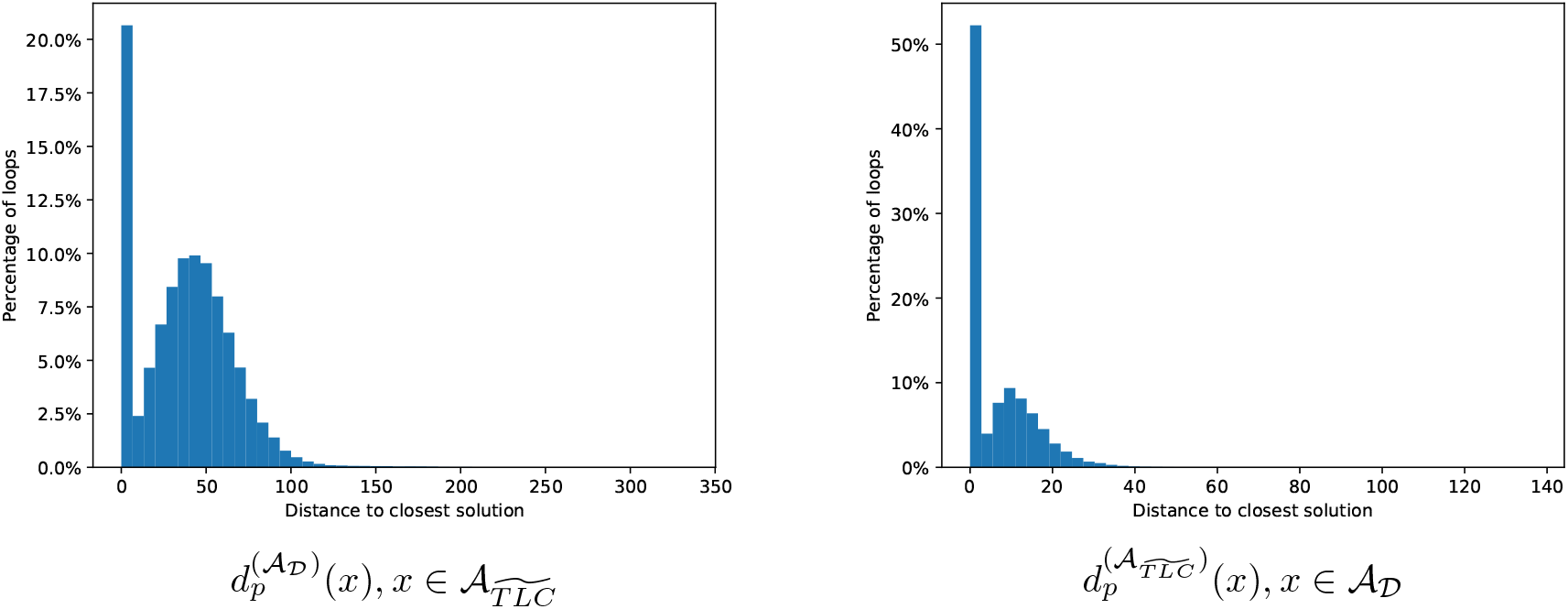
Distances to nearest neighbors, see Eq. 7, in degrees.

**Figure S 6:**
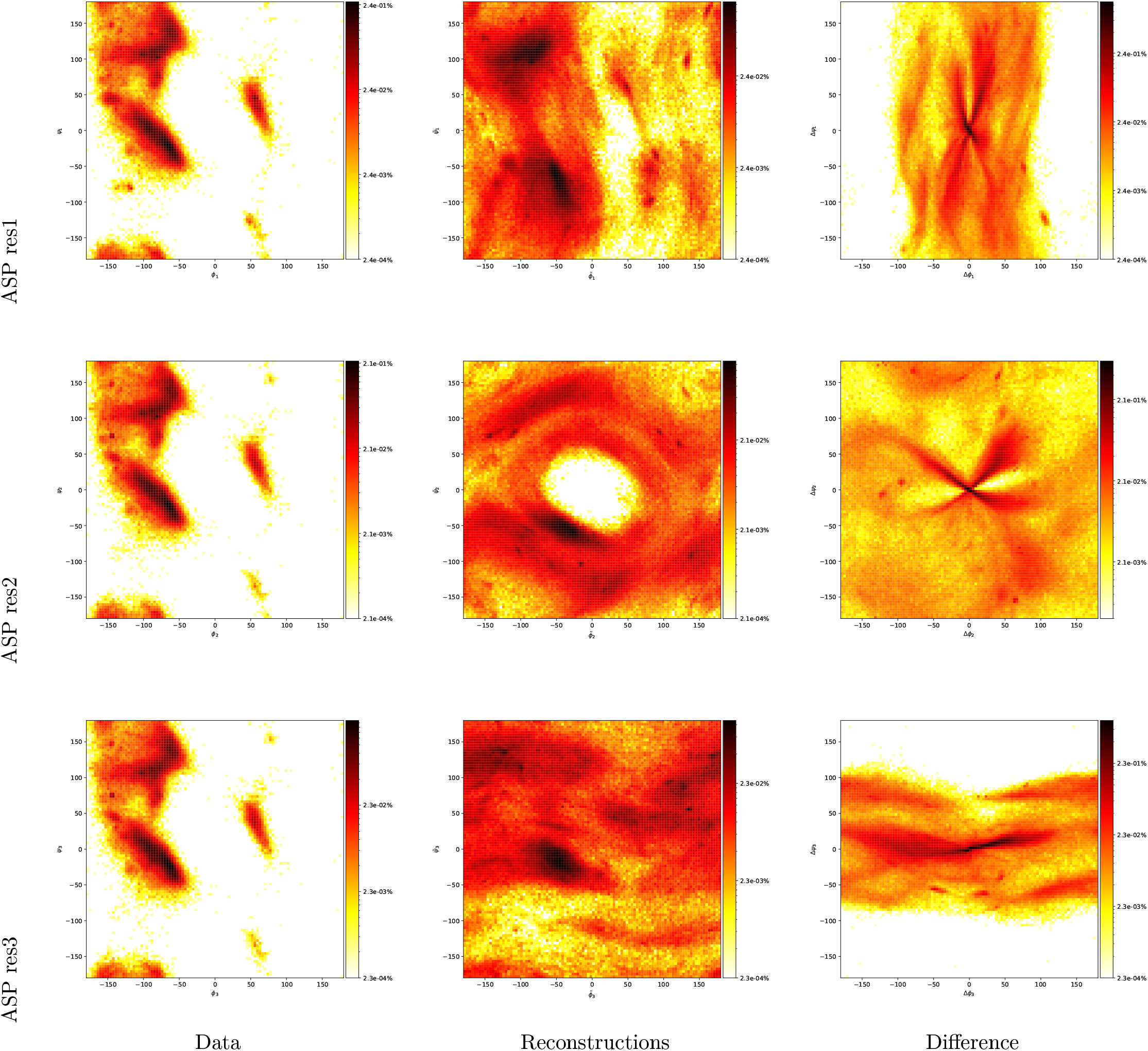
Amino acid: ASP. **(Left column)** Distributions in Ramachandran domains 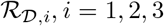, **(Middle column)** Distributions in Ramachandran domains 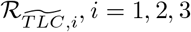 **(Right column)** Difference

**Figure S 7:**
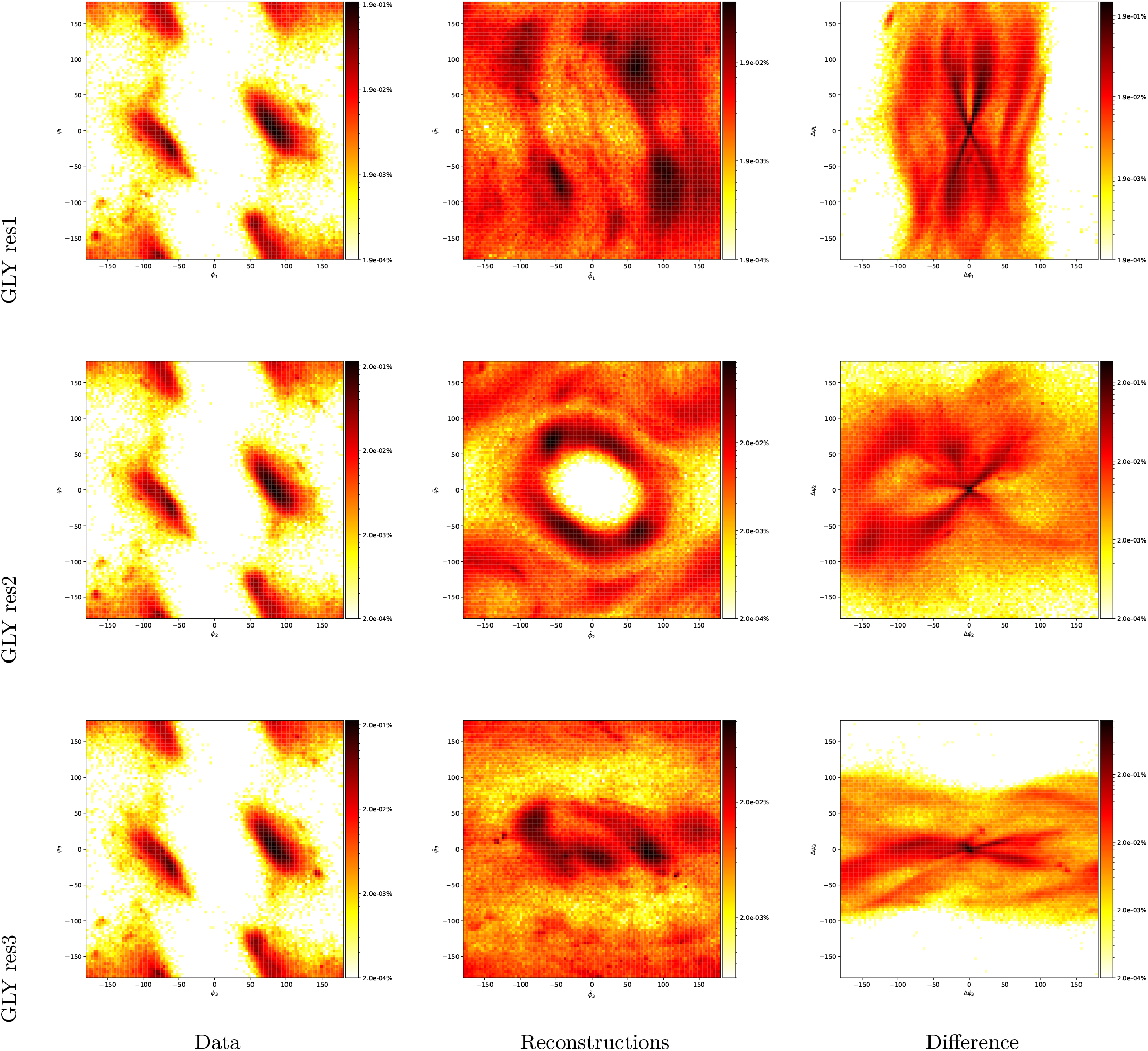
Amino acid: GLY. **(Left column)** Distributions in Ramachandran domains 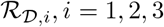, **(Middle column)** Distributions in Ramachandran domains 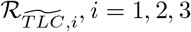 **(Right column)** Difference

**Figure S 8:**
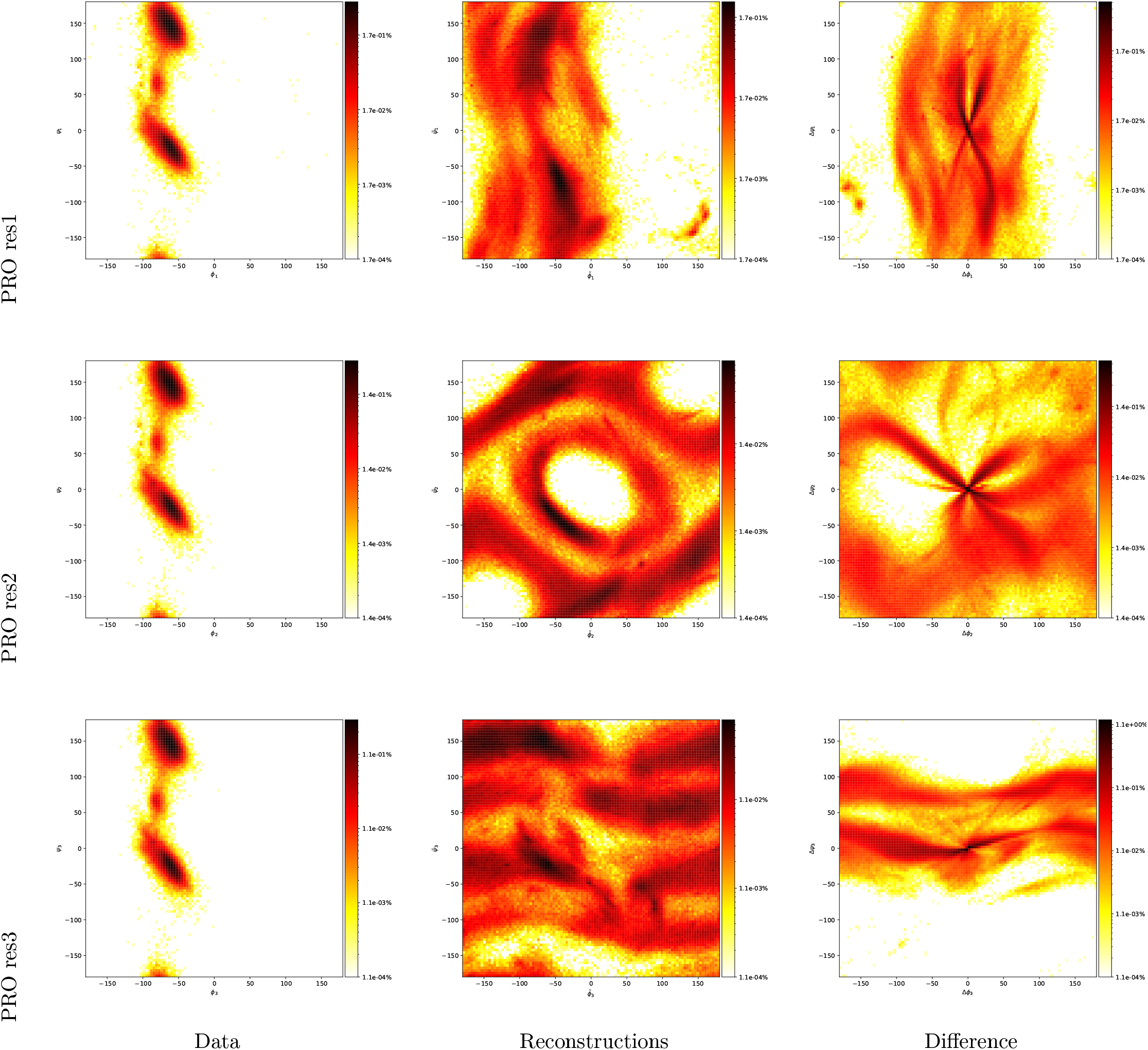
Amino acid: PRO. **(Left column)** Distributions in Ramachandran domains 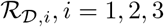, **(Middle column)** Distributions in Ramachandran domains 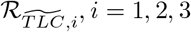 **(Right column)** Difference

**Table S 1:**
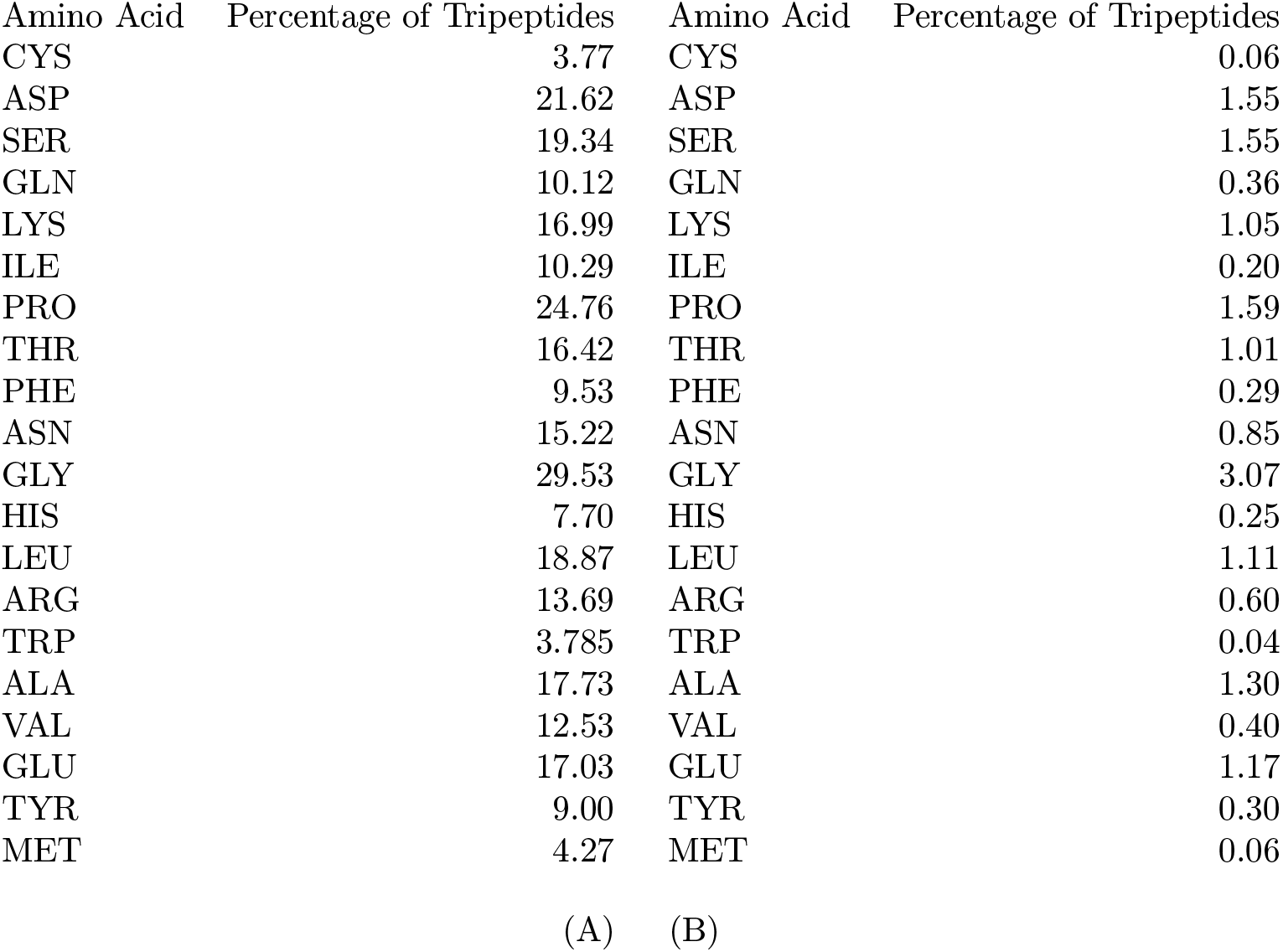
Amino acid composition of tripeptides. **(A)** Percentage of tripeptides containing the indicated amino acid at least once. **(B)** Percentage of tripeptides containing an amino acid at least twice.

**Table S 2:**
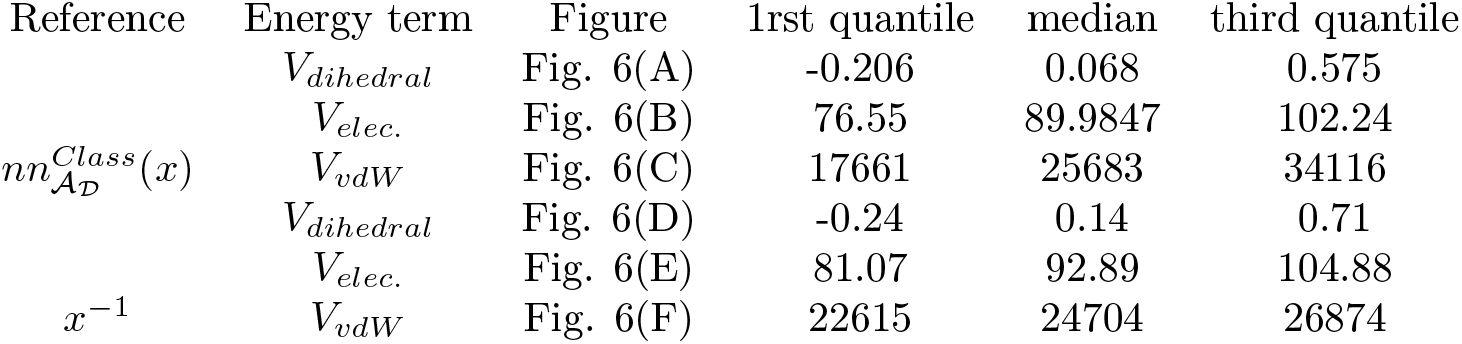
Table of Δ*V*_∗_ in kcal/mol.

**Table S 3:**
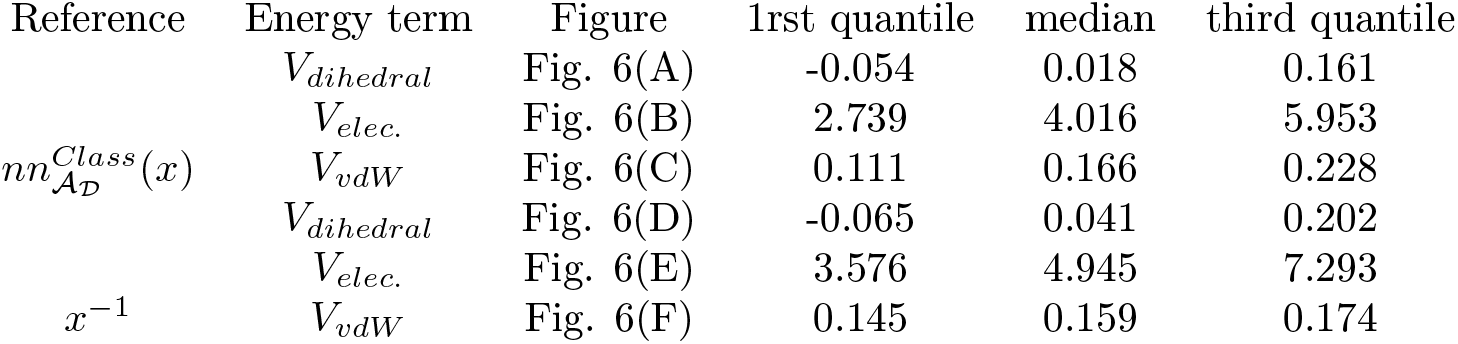
Table of Δ*_r_V*_∗_ ratios.

